# Event-Centered Prediction: How Future Interaction Points Shape Human Anticipation of Motion

**DOI:** 10.64898/2026.07.15.738481

**Authors:** Gonzalo Aparicio-Rodríguez, Tyssa Martín-Fernández, Paloma Manubens, Abel Sánchez-Jiménez, Carlos Calvo-Tapia, José Antonio Villacorta-Atienza

## Abstract

Prediction in dynamic situations, in which relevant elements evolve over time, is a fundamental cognitive function. The brain relies on specialized predictive mechanisms, including time compaction, a process that supports dynamic processing by embedding temporal information into space and transforming future interactions into salient spatial representations. Here we investigated how future interactions are salient during dynamic events and how this salience shapes behavior. Participants performed a visuomotor prediction task in which they estimated the future trajectory of a moving object after observing only the initial portion of its motion, while another object was simultaneously present and could generate either interactive (collision) or non-interactive (crossing) dynamics. Although accurate performance required extrapolating motion solely from kinematic information, participants’ predictions were systematically biased toward locations associated with future interactions. Prediction accuracy was reduced in situations involving potential future interactions compared to non-interactive dynamics. Importantly, participants consistently responded closer to predicted interaction points, even when this strategy did not improve accuracy or trajectory extrapolation. Substantial inter-individual variability was observed, revealing conservative and risk-taking predictive strategies with systematic group differences. When participants were explicitly instructed to improve performance, overall accuracy improved only marginally, while predictive behavior shifted toward greater reliance on interaction-related locations, particularly among those who had not already adopted this strategy. We propose that this interaction-driven bias reflects a core property of time compaction, supporting the idea that predictive cognition relies on future interactions as stable reference points under dynamic uncertainty.

## INTRODUCTION

Prediction is a core component of human behavior^1^ and operates simultaneously across multiple cognitive levels^2^. However, prediction does not function uniformly. Instead, it selectively prioritizes stimuli that are most relevant to the organism^3, 4^. This prioritization allows the brain to conserve resources while preparing the individual for appropriate responses^5^. Natural environments are inherently dynamic, involving interactions among multiple elements and the consequences that emerge from those interactions^6^. As a result, purely reactive cognition is insufficient: by the time a response is generated, the opportunity for effective action has often already passed. In such contexts, the isolated motion of environmental elements is not meaningful on its own; it becomes relevant only when considered within the full causal structure. Namely, the movements of the elements and the consequences those movements entail for themselves and for other interacting elements.

Prediction can be understood as a fundamental strategy for coping with the uncertainty inherent in complex systems, favoring the survival of individuals who are less constrained by the stochastic dynamics of their environment^4^. These stochastic dynamics manifest as uncertainty, a state in which an organism’s internal representations are unable to accurately forecast future events or environmental states^7^. The brain must therefore continuously manage uncertainty, relying on probabilistic mechanisms to anticipate upcoming events, meaning that when probability is high, estimation is precise^8, 9^, while also using uncertainty itself as a signal of the end of events in order to update internal representations^10, 11^.

Most research in this field has traditionally relied on static stimuli, even though natural environments are predominantly dynamic (that is, composed of elements in motion) and therefore inherently associated with greater uncertainty^12^. To cope with such dynamic environments, multiple brain regions anticipate aspects of motion before they occur, operating across different hierarchical levels of abstraction^12^. How does the brain accomplish this? A prominent account proposes that humans rely on an internal physics system capable of generating approximate representations of physical dynamics, allowing predictions about how objects will move and interact^13, 14^.

However, these internal physics models are not deterministic. Humans predict the outcomes of even simple object motions with a degree of uncertainty that cannot be explained solely by sensory noise^15^. The problem becomes even more pronounced when objects interact. Cause-effect interactions introduce additional uncertainty into the system, and predictive accuracy decreases as the number of interactions increases^15^. Complementary evidence further shows that increasing the number of causal relationships in a scenario recruits additional neural resources and leads to stronger activation in brain regions involved in planning and prospective reasoning^16^.

Taken together, these findings highlight interactions and causal relations as critical prediction horizons: temporal points beyond which uncertainty grows substantially, and the brain struggles to reliably anticipate how the situation will unfold. Because interactions are ubiquitous in natural environments, the brain must rely on strategies that allow it to cope with these prediction horizons and anticipate outcomes efficiently. Therefore, a future interaction is not merely a predictable point in time, but a moment at which a causal transition and a shift in system dynamics are expected to occur^6^. Examples of such interactions include collisions or contacts, which in biological contexts range from hunting prey to, in contemporary settings, navigating a crowded street. These interactions segment time by defining events with clear causal consequences, and their significance makes them key reference points in the temporal representation of the future.

Indeed, research has shown that a cognitive mechanism, known as time compaction, facilitates the processing of dynamic situations by reducing them to static representations of their interaction structure^17^. Prior evidence indicates that this compacted internal representation (CIR) of future interactions plays a critical role in both human memory^18^ and decision-making^19^, and compatible results of time compaction having a role in rat^20^ and bat^21^ perception have been found.

In the present work, we aim to unravel how the brain responds to uncertainty in prediction. To do this, we examine whether these future interaction points constitute regions of heightened subjective certainty in prediction, even when geometrically simpler predictive alternatives are available, such as internal physics-based models. We show that interaction points are salient when predicting the trajectory of a ball that collides with another ball. While women use more conservative approaches by predicting near the interaction point, men tend to take more risks further from this spot However, when asked to improve the accuracy of their predictions, men predict closer to the interaction point. We argue that human prediction is not merely a matter of physically extrapolating motion, but is instead organized around events, reflecting a predictive structure centered on future interactions.

## MATERIALS AND METHODS

### Experimental design

To assess how anticipated interactions shape predictive processing, participants completed a computerized task presenting brief dynamic scenes from an allocentric viewpoint. Each trial depicted two spheres, red and green, moving across a white field. After two seconds of motion, the display vanished, and participants indicated by clicking on the screen the location corresponding most closely to the unseen continuation of the red sphere’s trajectory.

Stimuli were categorized into two types based on the relative motion of the spheres: *cross* and *collision*. In cross trials, the trajectories intersected without physical interaction, whereas in collision trials, the spheres collided and altered their paths upon impact. Time compaction was therefore expected to occur only in collision trials, enabling assessment of how future interactions influence prediction. The point of contact between spheres in collision trials is referred to as the *collision point*; an analogous *crossing point* was defined in cross trials, corresponding to the spatial intersection of the trajectories.

To ensure correct identification of cross and collision trials, each stimulus type was presented in separate experimental phases. Because collision trials were expected to be more demanding, potentially leading participants to adopt more conservative predictive strategies, the cross phase was always administered first, followed by the collision phase, to minimize carryover effects. Both phases were repeated in a second session, identical in structure but accompanied by instructions to improve performance.

For each trial, two dependent measures were derived: error and distance (Fig. 1). Error corresponded to the minimal distance between the participant’s click and the true trajectory of the red sphere. Distance represented the minimal distance between the click and the crossing or collision point and could take positive or negative values depending on which side of a defined bisector line the response fell. The bisector was formed by (i) the vectors of the green sphere’s trajectory before and the red sphere’s trajectory after the intersection in cross trials, or (ii) the red sphere’s vectors before and after the collision in collision trials. Responses located on the side preceding the crossing or collision were assigned negative values; those beyond the interaction point were assigned positive values. All measures were expressed in screen-based units to control for potential hardware variability.

**Figure 1.**
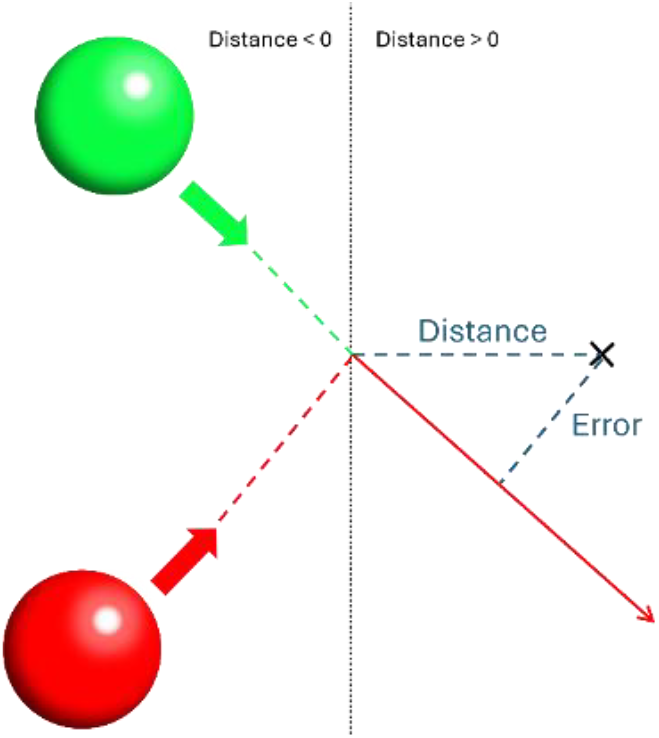
Schematic representation of the measures obtained in each trial. The figure illustrates a collision between the red and green spheres. Solid arrows indicate the visible trajectories presented to participants, while dashed lines represent the unseen continuations prior to or following the collision. The participant’s response is shown as a black “×”. *Error* corresponds to the shortest distance between the click and the red sphere’s trajectory. *Distance* is defined as the shortest distance from the click to the collision point. The black dashed line denotes the bisector formed by the red sphere’s trajectory vectors before and after the collision. Clicks falling on the pre-collision side of the bisector yield negative distance values, whereas those on the post-collision side yield positive values.

### Experimental procedure

The task was implemented in C++ and JavaScript. Red and green spheres were rendered with shading to enhance three-dimensionality, and all spatial parameters scaled with screen size. The interface maintained a 2:1 aspect ratio, corresponding to an internal coordinate system of 8 × 4 units, which defined object dimensions and reference points (e.g., start, disappearance, and interaction locations). Each sphere had a fixed diameter of 0.2 units. Distinct cross and collision stimuli were generated by varying the spheres’ initial positions, motion vectors, and velocities.

Each phase comprised 16 unique trials. Participants were informed of the phase type (collision or cross) via on-screen text and an auditory cue. The first three trials served as practice, after which the red sphere’s full trajectory and the participant’s response point were displayed to ensure task comprehension. Trial order was identical across participants. Following stimulus disappearance, participants had two seconds to indicate their response; omissions were recorded as null trials. Auditory cues signaled response timing, and timeouts were indicated by both a visual message and an alarm sound.

Before testing, participants viewed an instructional video outlining the task. They were informed that each trial would display two moving spheres that would disappear before crossing or colliding, and that their task was to click on the predicted continuation of the red sphere’s trajectory within two seconds. The video also explained the session structure (cross trials first, collision trials second) with three initial practice trials per session and illustrated examples of the full stimuli. Participants were instructed not to trace the motion with their hands or other objects and were informed that performance feedback would be provided after each trial. Any remaining questions were resolved by referring to the video instructions.

The test was performed in calm environments individually or in groups, but in the latter case on individual computers avoiding all sorts of interaction among participants.

### Participants

104 people (64 female and 40 male) completed the whole test, including both sessions. Age of participants ranged from 18 to 27 years old, with similar age distributions for men and women (Table S1). All of them provided informed consent to anonymously take part in the experiments, according to the experimental procedures approved by the Institutional Review Board (Committee of Bioethics, National Distance Education University). All experiments were performed following the guidelines and regulations set forth by the Declaration of Helsinki.

### Statistical analysis

Statistical analysis was performed in R software (v. 4.2.1).

Each participant’s response to a given stimulus was treated as an individual observation. Trials with response times below 300 ms were excluded, as such latencies are incompatible with deliberate decision-making and likely reflect random responses^22–24^.

Error was analyzed using linear mixed-effects models (*lme4*^*25*^), with significance values estimated via *lmerTest*^26^. Models included participant ID and stimulus identity as random effects, and session, stimulus type (collision or cross), and gender as fixed factors. Predictor selection followed a backward stepwise elimination procedure. To satisfy normality assumptions, error values were Box–Cox transformed^27^ for inferential testing, while effect sizes were derived from models fitted to untransformed data.

Distance was first analyzed by estimating the probability of clicks across distance bins (0.2-unit intervals), computed separately by session, gender, and stimulus type. Resulting distributions were compared using a two-sample test based on the DTS statistic (*twoSamples* package^28^).

To further examine the influence of the collision point on predictive responses, we conducted an areal analysis defining four spatial regions relative to the crossing/collision point (Fig. 2). Zone B corresponded to the collision/crossing zone, defined as a circle centered on the interaction point with a radius of 0.22 units (110% of the sphere diameter) to fully encompass both spheres at contact. Zone A represented the disappearance zone: a semi-annulus contiguous to Zone B on the negative-distance side with a width of 0.376 units, calculated by subtracting the radius of Zone B from the mean distance between the disappearance point and the crossing/collision point. Zones *C* and *D* served as control regions, defined as semi-annuli on the positive-distance side with widths of 0.44 units. Zone C was adjacent to Zone B (inner radius = 0.22 units), whereas Zone D was separated from Zone C by 0.22 units (inner radius = 0.88 units). Zone D was used as a reference for tracking distant clicks.

**Figure 2.**
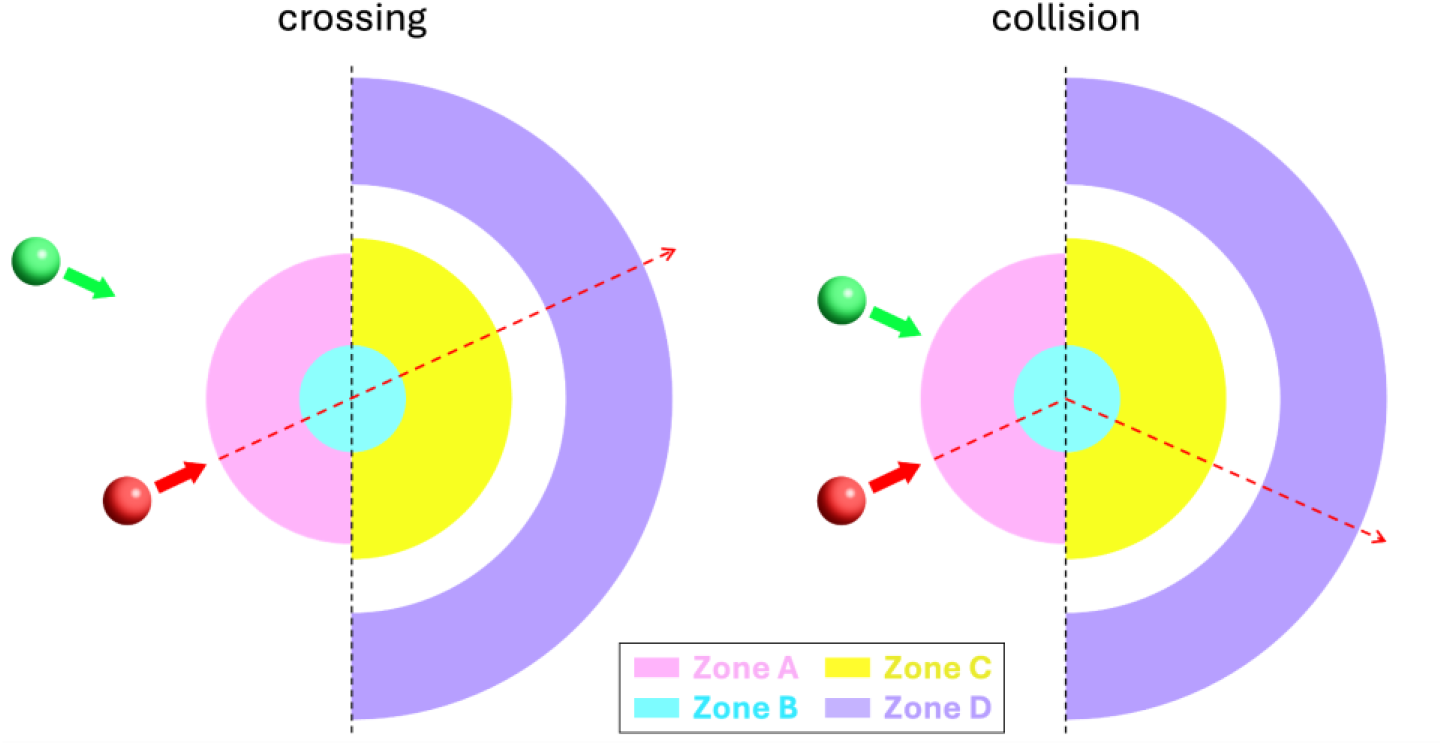
Scaled diagram of the zones established for statistical analysis. Left panel depicts the zones for a crossing stimulus and right panel depicts the zones for a collision stimulus. Solid thick arrows represent the balls’ direction and red dashed arrows represent the trajectory of the red ball once the stimuli had disappeared. The dashed black line marks the bisect line to the vector of the balls from which distances change from negative to positive (see Fig. 1). The zones are labeled in pink for Zone A, blue for Zone B, yellow for Zone C, and purple for Zone D. All zones are centered on the crossing/collision point. Zone A is a semi-annulus that only admits negative distance values between 0.376 units and 0.22 units. Zone B is a circle of radius = 0.22 units. Zone C is a semi-annulus that only admits positive distance values of width = 0.44 units and radius of the internal circle = 0.22 units. Zone D is a semi-annulus that only admits positive distance values of width = 0.44 units and radius of the internal circle = 0.88 units.

To analyze click frequency across zones, each observation was replicated four times (once per zone), and a binary variable was created to indicate whether a click occurred within each zone (1) or not (0). This binary outcome was modeled using a generalized linear mixed-effects model (*lme4*^25^) with a logit link and participant ID as a random effect. Stimulus identity was excluded, as its inclusion did not capture additional variance. The initial fixed factors were gender, zone, and stimulus type, with predictor selection based on backward stepwise elimination. We used anova tests to assess significance for triple interaction models involving categorical variables. Effect sizes were derived using the *ggeffect* function (*ggeffects*^29^), and significance of pairwise contrasts was assessed via estimated marginal means (*emmeans*^30^).

## RESULTS

All 104 participants completed both experimental sessions. Error in prediction was significantly lower for cross than for collision stimuli, both for Session 1 (Fig. S1A; *p* < 0.001; Table S2) and for Session 2 (Fig. S1B; *p* < 0.001; Table S3). Although overall accuracy slightly improved in Session 2 (*p* < 0.001; Table S4), the difference between stimulus types remained, and the magnitude of improvement was negligible (≈0.02 spatial units; Table S5). Thus, cross trajectories were consistently easier to predict across sessions.

### Distance distributions and saliency of the interaction point (Session 1)

Click distributions as a function of distance differed significantly between cross and collision stimuli, for both women (Figs. 3A; *p* < 0.001; Table S6) and men (Figs. 3B; *p* = 0.002; Table S6). For cross trials, men and women exhibited similar (*p* = 0.107; Table S6) normal-like distributions centered away from the red ball’s disappearance point, indicating that participants did not adopt a trivial conservative strategy and instead tolerated greater predictive uncertainty although, as already mentioned, this risk-taking behavior was associated with higher overall accuracy.

**Figure 3.**
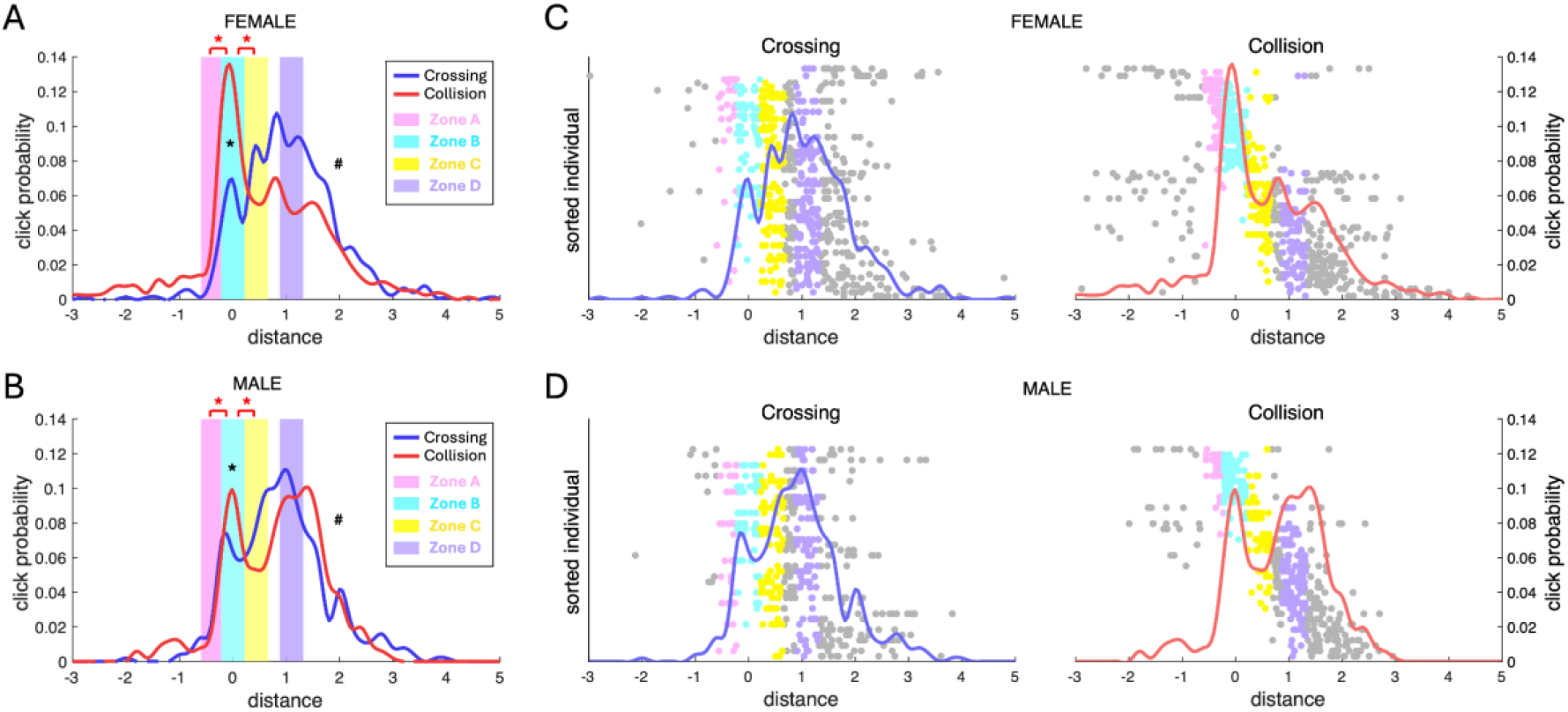
Click distributions for Session 1. **A.** Click distributions for female participants in Session 1. Blue and red curves represent the proportion of clicks within 0.2-unit distance bins for cross and collision stimuli, respectively. Shaded regions indicate the spatial areas defined in Fig. 2. Hashtags denote significant differences between cross and collision distributions; black asterisks mark significant differences in click frequency within Zone B; red asterisks indicate significant differences between zones. **B**. Click distributions for male participants in Session 1. Same conventions as in A. **C**. Individual click patterns for female participants. Each dot represents a single click, plotted by distance (x-axis) and participant (y-axis). Participants are ordered by their mean click distance for collision stimuli, with the shortest mean distance at the top. Dot color indicates the corresponding zone (Fig. 2), while grey dots indicate clicks outside defined zones. The left panel shows cross trials (blue), and the right panel shows collision trials (red), with the corresponding aggregate distributions, represented by the colored function. **D**. Individual click patterns for male participants. Same conventions as in C.

In contrast, collision trials showed an additional accumulation of clicks near the collision point, indicating that this location exerted a strong attractive effect on predictions. This tendency was more pronounced in women than in men, showing gender-differenced distributions (*p* < 0.001; Table S6). Nevertheless, for both genders, Zone B (collision area) was clicked significantly more often in collision than in cross trials, despite the existence of an analogous spatial point in the latter (Fig. 3A and B; Fig. S2; *pwomen* < 0.001; Tables S7-10; *pmen* = 0.015; Tables S7, 8, 12, and 13). Moreover, in collision trials only, Zone B was significantly more likely to be selected than its adjacent regions (Zones A and C) (*pwomen B vs A* < 0.001 and *pwomen B vs C* < 0.001; Tables S7-9 and 11; *pmen B vs A* < 0.001 and *pmen B vs C* = 0.004; Tables S7, 8, 12, and 14).

Substantial inter-individual variability was observed (Figs. 3C and D; random-intercept variance = 0.32; Table S7), particularly for collision stimuli. While responses to cross stimuli were relatively homogeneous across participants, collision trials revealed participants tended to use either one out of two of the following approaches to the task: clicking near the collision point or responding at larger distances. This pattern indicates that future interactions induce a conservative strategy in only a subset of participants. Notably, this conservative strategy did not involve approaching the disappearance point (which would indeed be the easier approach) but rather targeting the collision point, highlighting its saliency. Gender differences reflected differences in subpopulation composition, with women showing a higher proportion of conservative responders and men a higher proportion of risk-takers. Stimulus identity did not explain significant variance beyond participant-level effects (Fig. S3).

Overall, these findings demonstrate that anticipated future interactions render the collision point salient, acting as an attractor for predictive responses even when this strategy does not maximize accuracy.

### Effect of instruction (Session 2)

In Session 2, participants were instructed to improve performance. For cross stimuli, distance distributions changed significantly relative to Session 1 for women (Fig. 4A, left panel; *p* < 0.001; Table S15) and men (Fig. 4B, left panel; *p* = 0.007; Table S15), with clicks clustering closer to the disappearance point, reflecting a more conservative shift. For collision stimuli, no session-related changes were observed in women(Fig. 4A, right panel; *p* = 0.661; Table S15); however, men showed a significant shift toward the collision point in Session 2 (Fig. 4B, right panel; *p* = 0.003; Table S15), yielding a resembling distribution to those observed in women across both sessions.

**Figure 4.**
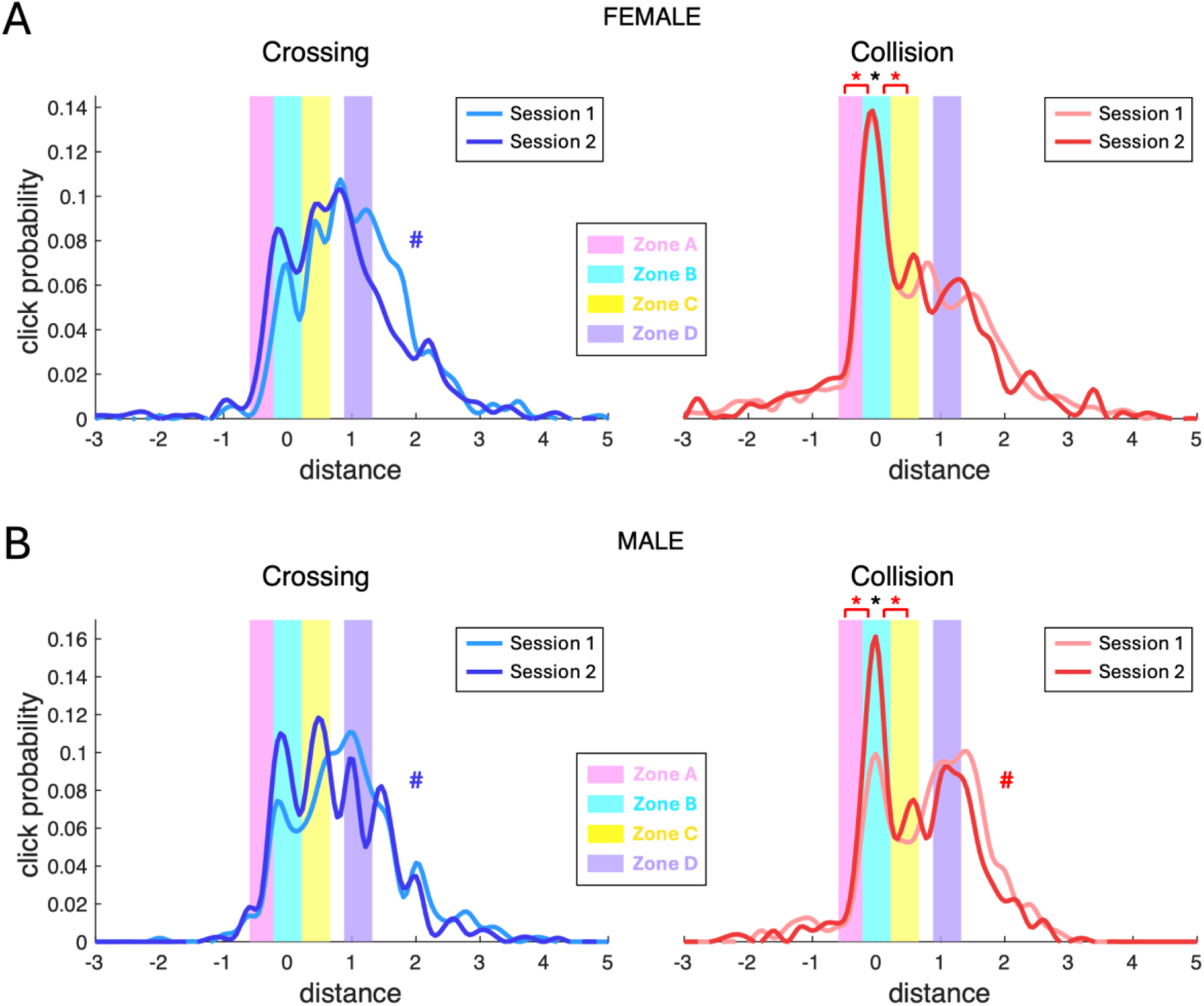
Changes in click distributions between sessions. **A.** Differences in click distributions for female participants. Light and dark blue curves indicate the proportion of clicks in 0.2-unit distance bins for cross stimuli in Sessions 1 and 2, respectively; light and dark red curves indicate the same for collision stimuli. Shaded regions correspond to the spatial areas defined in Fig. 2. Hashtags denote significant differences between Session 1 and Session 2 distributions for each stimulus type; black asterisks indicate significant differences in click frequency within Area B between collision and cross stimuli in Session 2; red asterisks indicate significant differences between areas for collision stimuli in Session 2. **B**. Differences in click distributions for male participants. Same conventions as in A.

The saliency of the collision point persisted in Session 2 for both genders. In collision trials, Zone B was clicked significantly more often than in cross trials (Fig. 4; Fig. S4; *pwomen* < 0.001; Tables S16-19; *pmen* = 0.001; Tables S16, 17, 21, and 22) and more frequently than adjacent Zones A and C (*pwomen B vs A* < 0.001 and *pwomen B vs C* < 0.001; Tables S16-18 and 20; *pmen B vs A* < 0.001 and *pmen B vs C* < 0.001; Tables S16, 17, 21 and 23).

These session effects were again driven by inter-individual differences (random-intercept variance = 0.26; Table S16). The shifts observed for this session in this individual trends are different. While for cross trials, participants tended to click at shorter distances in Session 2 (Fig. 5), for collision trials participants who initially adopted riskier strategies in Session 1 adjusted by clicking closer to the collision point in Session 2 (Fig. 6). As in Session 1, stimulus identity did not account for significant variance in click distance (Fig. S5).

**Figure 5.**
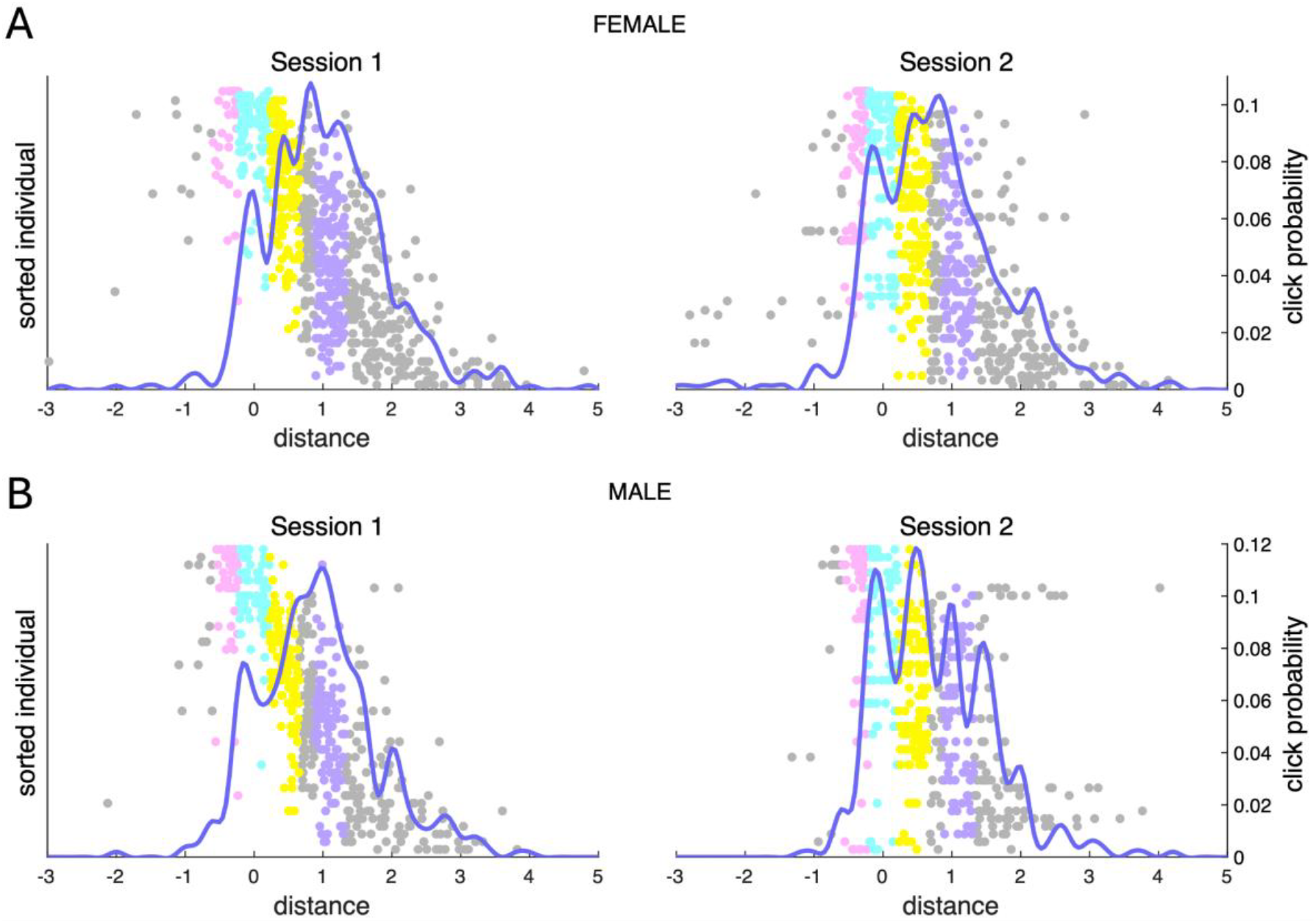
Change in click strategy for cross stimuli between sessions. **A.** Change in click strategy for cross stimuli between sessions for female participants. Each dot represents a single click, plotted by distance (x-axis) and participant (y-axis). Participants are ordered by their mean click distance for cross stimuli in Session 1, with the shortest mean distance at the top. Dot color indicates the corresponding zone (Fig. 2), while grey dots indicate clicks outside defined zones. The left panel shows clicks in Session 1, and the right panel shows clicks in Session 2, with the corresponding aggregate distributions, represented by the colored function, also used in Fig. 4. **B**. Change in click strategy for cross stimuli between sessions for male participants. Same conventions as in C.

**Fig 6.**
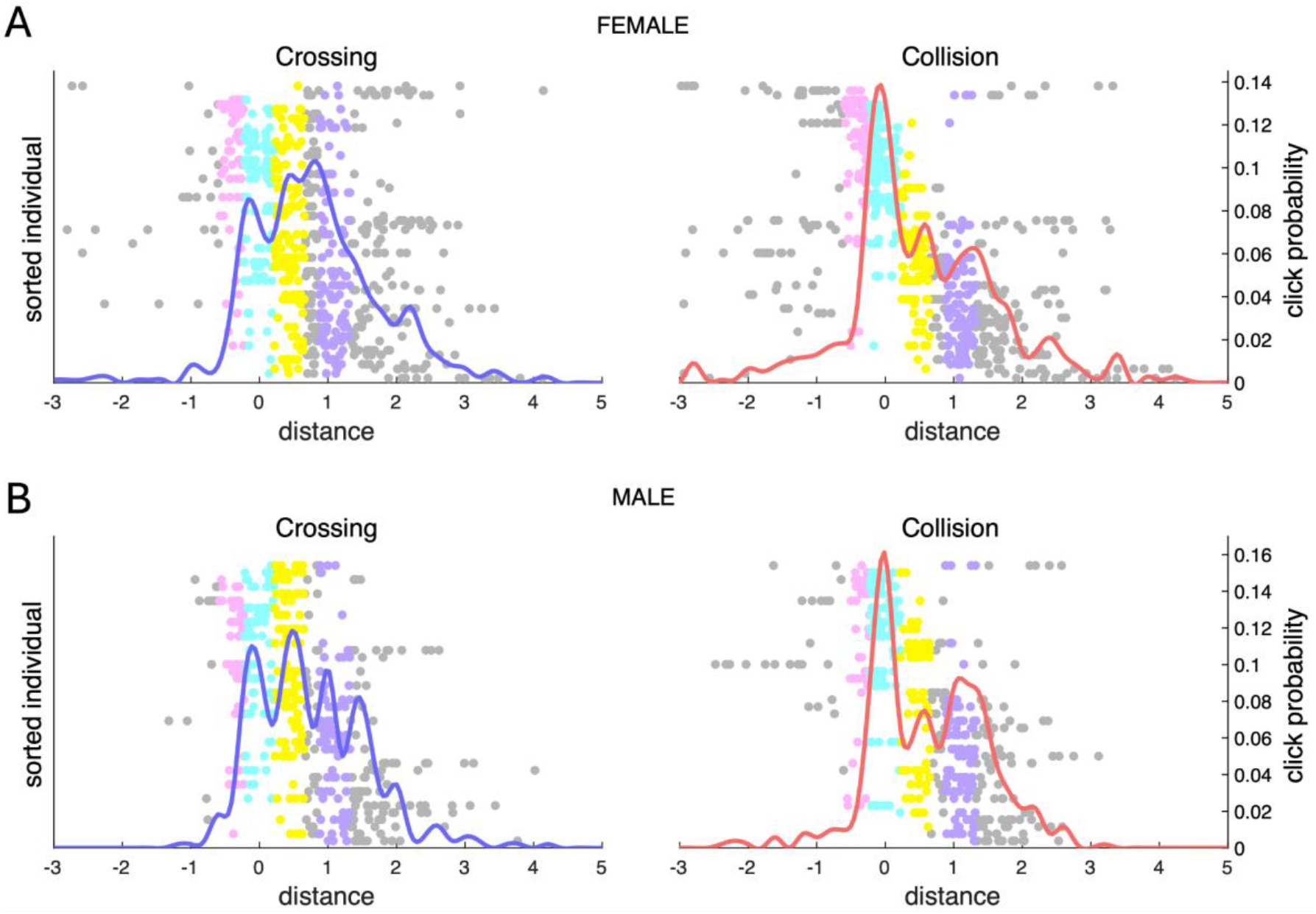
Individual click patterns in Session 2. **A**. Individual click patterns for female participants. Each dot represents a single click, plotted by distance (x-axis) and participant (y-axis). Participants are ordered by their mean click distance for collision stimuli in Session 1, with the shortest mean distance at the top. Dot color indicates the corresponding zone (Fig. 2), while grey dots indicate clicks outside defined zones. The left panel shows cross trials (blue), and the right panel shows collision trials (red), with the corresponding aggregate distributions, represented by the colored function, also used in Fig. 4. **B**. Individual click patterns for male participants. Same conventions as in C.

Together, these results indicate that while participants can adopt more conservative strategies when instructed to get better results, the nature of such strategies depends on stimulus structure. For cross trials, conservatism involves clicking at shorter distances, whereas for collision trials it entails targeting the anticipated interaction point. Female participants already exhibited this latter strategy in Session 1, explaining the absence of further session-related changes.

## DISCUSSION

This work investigates human prediction in dynamic stimuli involving future interactions and shows that, despite marked intra-individual variability, participants tend to shift their responses toward future interaction points when aiming to improve accuracy, even though this strategy does not necessarily enhance performance. This pattern suggests that prediction in dynamic contexts does not unfold linearly across the entire future trajectory but is instead biased toward events involving causal transitions. Thus, although the human brain is equipped with an internal physics system capable of predicting complex dynamic outcomes^14^, future interaction points appear to serve as anchors for prediction.

These findings are consistent with time compaction, which proposes that dynamic situations are represented as static maps of future interactions, effectively reducing the temporal dimension to a spatial structure centered on salient interaction points^17, 31^. If prediction operates on such compacted representations, these points would naturally generate a sense of subjective certainty. Crucially, geometric extrapolation, based on internal physics models, was available in our task; therefore, the observed preference for interaction points cannot be attributed to the absence of alternative strategies. Even when a theoretically simpler and potentially more accurate response was possible, participants privileged locations around behaviorally relevant events. Prediction thus appears to be organized around events rather than unfolding as a continuous dynamic process, with time compaction functioning as an organizing principle that constrains how future states are represented. Note that time compaction itself relies on an initial process of linear geometric prediction to estimate future interaction locations^17^, suggesting that linear geometric and event-centered prediction operate concurrently.

More broadly, these results highlight that interaction points are not merely spatial coordinates where events occur, but loci of semantic and behavioral significance^32^. An interaction point may represent a goal to be reached (e.g., catching up with someone we want to talk to who is walking ahead), a danger to be avoided (e.g., sidestepping a bicycle), or a location at which an action must be executed (e.g., taking a plate from a catering waiter). This framework also extends to higher-order causal chains, in which the interaction between A and B triggers a subsequent interaction between B and C, as in a billiard table scenario. Time compaction provides an efficient and low-cost solution for representing and navigating such dynamically structured environments.

Moreover, our results raise the question of whether saliency of future interactions in prediction processes is a consequence of the use of time compaction or rather the preference of interaction points for prediction is the root of time compaction. The first case would be a straightforward extension of the already known role of time compaction in human behavior, as future interaction points are also salient in memory^18^ and decision-making^19^. On the other hand, the salience of future interactions might also be the evolutive origin of time compaction. It is thought that prediction lies in the core of cognition^1^, supporting representation of dynamic events, which include anticipated effects of action^33^. Anticipation to future events allows denoising perceptive streams, providing stable and coherent representations^34^. This way, prediction gives rise to latent representations that guide behavior^35^. In 2021, Recanatesi and colleagues^36^ showed that predictive coding naturally leads to compaction of representations (*i*.*e*. reducing dimensionality of the latent state), by detecting the regularities of the situation, in a similar way as proposed by time compaction^17^. It is well known that when obtaining feedback in uncertainty tasks, internal representations are updated to reflect lack of predictive certainty^7, 37, 38^ and only representations that yield successful behaviors would be stored in memory^39^, keeping information about the conditions in which the representation rose and its utility^40^. Thus, following this hypothesis, representations of dynamic situations would be updated and then stored during learning closer to the interaction point, the latest spatiotemporal point of prediction certainty (as mentioned in the introduction, a prediction horizon). Moreover, danger and benefits in dynamic situations take place in the interaction point (for instance imagine a predator haunting its prey), while posterior states to these interactions are often irrelevant. Briefly, the relevance of these spots and core predictive coding processes might have given birth to time compaction by naturally providing a solution to the information load of dynamic situations and their inherent uncertainty through low-dimensional representations. While our present results do not allow us to unravel this question, we expect upcoming research to throw some light regarding it. Although our results do not clarify this cause–effect relationship, they do show that prediction organized around events is behaviorally functional.

Either case, under the time compaction framework, how can the salience of future interactions when predicting be explained at a circuit level? In humans, prediction is universal: multiple cognitive functions are devoted to predictive processes and almost every brain region participates in them^41^. Time compaction is also hypothesized to be ubiquitous, as it is thought to participate in many cognitive domains^42^ (*e*.*g*., Theory of Mind^43^) but also anatomically, as a mechanism as complex and widely used such as time compaction must be allocated in multiple brain structures. Therefore, time compaction might be favoring the salience of impending interactions and several levels and through several neuronal circuitries. However, we propose hereunder an illustrative example on how time compaction might affect prediction in the hippocampal complex, a region that has already been thought to host part of the time compaction process^18, 20, 44^.

In a previous work^18^, we showed that stimuli containing future interactions are better memorized and recalled than stimuli not containing future interactions and we proposed that the reason of this was rooted in the concept of engram, as time compaction would allow for allocating the engram in hippocampal simpler circuits, therefore being more resistant to interferences and forgetting. Indeed, hippocampal complex has been shown to participate in prediction^45–48^ and it has been shown that memory and prediction are linked, as more predictable stimuli are also better remembered^49^. Particularly, once a sequence has been memorized, when the onset of the sequence is perceived, its activation pattern is similar to activation pattern of the remainder of the sequence^50^, so the onset works as a cue of the hippocampal representation of the remainder^51^. In this manner, we propose that the same circuits responsible for memorizing compacted stimuli are also responsible for predicting this kind of stimuli. Following time compaction, these situations are represented by a single point in space, the interaction point^17^, the same point that would be codified by the corresponding engram^18^. Therefore, when aiming to better predict the collision stimuli from our experiment, interaction points are perceived as more accurate spots for prediction, due to the constraints of the circuit, which is focused on representing and memorizing the interaction point. On the other hand, as crossing points are not represented (due to the lack of interaction), this spot does not seem to provide a safer prediction for participants in crossing stimuli.

Our results also reveal gender differences: women tended to gravitate toward the interaction point spontaneously, whereas men did so primarily when explicitly instructed to improve their accuracy. This convergence suggests that the interaction point is not a fixed attractor that rigidly determines behavior; rather, it appears to be a strategic resource that can be recruited when task demands require it. Prior time compaction research has also found gender differences in time compaction recurrence^18, 19^, but while in these past works men were the ones that under no-matter which context had a saliency of future interactions, our present results show the opposite situation. However, asking participants to try to overcome their previous results render men more conservative, resorting to time compaction in a more similar way women initially did. This is congruent with the existing literature, as men have been reported to be more prone to take risks than women^52, 53^ and tend to overestimate their own performance^54^, which would also explain the shift in strategy from phase one to phase two, as phase two task implicitly told participants that they had to reformulate the estimation of their performance and thus their strategy. While women were already predicting close to the location with greater certainty and could not improve much more their performance, men already could, yielding the shift in results.

Anyway, our results, together with previous experimental findings that showed that gender differences disappeared with increasing task complexity^18^, suggest that gender differences are due in contextual factors, which is something that has already been noticed by other researchers on the field of cognitive sciences^55, 56^. Moreover, unlike the other cases, here our experiment shows that these differences are not binary, but rather spectral, as individual strategies capture part of the variability of responses. These results are in line with recent insights on psychometrics that point that previously conceived binary differences in the field are indeed spectral^57^.

In summary, this work provides evidence of the salience of interaction points between moving elements when human participants aim to predict trajectories. As natural environments are inherently dynamic and the human brain works as a predictive machine^1^, this salience might be key to understanding human behavior and its neural correlates, as well as their evolutive history. Although future research is needed to interpret if this salience is the root of time compaction, this cognitive mechanism provides a framework to understand the role of future interactions in human cognition^42^.

## Acknowledgments

This research was supported by the National Project of the Ministry of Science, Innovation and Universities (MICIU/AEI), Spain, and by ERDF, EU, Grant PID2022-138659NB-I00, Spain, the European Social Fund Plus (ESF+), Grant PEJ-2024-AI/SAL-GL-32302 and pre-doctoral fellows from the Universidad Complutense de Madrid - Banco Santander (CT22/25).

## REFERENCES

1. Pezzulo G, Hoffmann J, Falcone R (2007) Anticipation and anticipatory behavior. Cogn Process 8:67–70

2. Ritter W, Sussman E, Deacon D, Cowan N, Vaughan HG (1999) Two cognitive systems simultaneously prepared for opposite events. Psychophysiology 36:S0048577299990248

3. Summerfield C, Egner T (2009) Expectation (and attention) in visual cognition. Trends Cogn Sci 13:403–409

4. Friston K (2010) The free-energy principle: a unified brain theory? Nat Rev Neurosci 11:127–138

5. Llinas RR (2001) I of the vortex: from neurons to self, Second printing.

6. Gibson JJ (1979) The ecological approach to visual perception. Houghton, Mifflin and Company, Boston, MA, US

7. Mushtaq F, Bland AR, Schaefer A (2011) Uncertainty and Cognitive Control. Front Psychol. 10.3389/fpsyg.2011.00249

8. Grabenhorst M, Poeppel D, Michalareas G (2025) Neural signatures of temporal anticipation in human cortex represent event probability density. Nat Commun 16:2602

9. Nobre AC, van Ede F (2018) Anticipated moments: temporal structure in attention. Nat Rev Neurosci 19:34–48

10. Baldwin DA, Kosie JE (2021) How Does the Mind Render Streaming Experience as Events? Top Cogn Sci 13:79–105

11. Nguyen TT, Etzel JA, Bezdek MA, Zacks JM (2026) Multiple event segmentation mechanisms in the human brain. 10.7554/eLife.107955.2

12. de Vries IEJ, Wurm MF (2023) Predictive neural representations of naturalistic dynamic input. Nat Commun 14:3858

13. McIntyre J, Zago M, Berthoz A, Lacquaniti F (2001) Does the brain model Newton’s laws? Nat Neurosci 4:693–694

14. Battaglia P, Ullman T, Tenenbaum J, Sanborn A, Forbus K, Gerstenberg T, Lagnado D (2012) Computational Models of Intuitive Physics. Proceedings of the Annual Meeting of the Cognitive Science Society 32–33

15. Smith KA, Vul E (2013) Sources of Uncertainty in Intuitive Physics. Top Cogn Sci 5:185–199

16. Kaplan R, King J, Koster R, Penny WD, Burgess N, Friston KJ (2017) The Neural Representation of Prospective Choice during Spatial Planning and Decisions. PLoS Biol 15:e1002588

17. Villacorta-Atienza JA, Velarde MG, Makarov VA (2010) Compact internal representation of dynamic situations: neural network implementing the causality principle. Biol Cybern 103:285–297

18. Aparicio-Rodríguez G, Ruiz-Navalón D, Manubens P, Sánchez-Jiménez A, Calvo-Tapia C, Villacorta-Atienza JA (2025) Efficient memorization of dynamic stimuli with future interactions. bioRxiv. 10.1101/2025.09.28.678599

19. Villacorta-Atienza JA, Calvo Tapia C, Díez-Hermano S, Sánchez-Jiménez A, Lobov S, Krilova N, Murciano A, López-Tolsa GE, Pellón R, Makarov VA (2021) Static internal representation of dynamic situations reveals time compaction in human cognition. J Adv Res 28:111–125

20. Dvorakova T, Lobellova V, Manubens P, Sanchez-Jimenez A, Villacorta-Atienza JA, Stuchlik A, Levcik D (2025) Spatial prediction of dynamic interactions in rats. PLoS One 20:e0319101

21. Dotson NM, Yartsev MM (2021) Nonlocal spatiotemporal representation in the hippocampus of freely flying bats. Science (1979) 373:242–247

22. Luce RD (1991) Response Times. 10.1093/acprof:oso/9780195070019.001.0001

23. Forstmann BU, Ratcliff R, Wagenmakers E-J (2016) Sequential Sampling Models in Cognitive Neuroscience: Advantages, Applications, and Extensions. Annu Rev Psychol 67:641–666

24. Ratcliff R, McKoon G (2008) The Diffusion Decision Model: Theory and Data for Two-Choice Decision Tasks. Neural Comput 20:873–922

25. Bates D, Maechler M, Bolker B, Walker S, Christensen RHB, Singmann H, Dai B, Scheipl F, Grothendieck G, Green P (2009) Package ‘lme4.’ URL http://lme4.r-forge.r-project.org

26. Kuznetsova A, Brockhoff PB, Christensen RHB (2015) Package ‘lmertest.’ R package version 2:734

27. Box GEP, Cox DR (1964) An Analysis of Transformations. J R Stat Soc Series B Stat Methodol 26:211–243

28. Dowd C (2020) A New ECDF Two-Sample Test Statistic.

29. Lüdecke D (2018) ggeffects: Tidy Data Frames of Marginal Effects from Regression Models. J Open Source Softw 3:772

30. Lenth R V. (2017) emmeans: Estimated Marginal Means, aka Least-Squares Means. CRAN: Contributed Packages. 10.32614/CRAN.package.emmeans

31. Villacorta-Atienza JA, Makarov VA (2013) Neural Network Architecture for Cognitive Navigation in Dynamic Environments. IEEE Trans Neural Netw Learn Syst 24:2075–2087

32. Calvo Tapia C, Villacorta-Atienza JA, Díez-Hermano S, Khoruzhko M, Lobov S, Potapov I, Sánchez-Jiménez A, Makarov VA (2020) Semantic Knowledge Representation for Strategic Interactions in Dynamic Situations. Front Neurorobot. 10.3389/fnbot.2020.00004

33. Schütz-Bosbach S, Prinz W (2007) Prospective coding in event representation. Cogn Process 8:93–102

34. Kveraga K, Boshyan J, Bar M (2007) Magnocellular Projections as the Trigger of Top-Down Facilitation in Recognition. The Journal of Neuroscience 27:13232–13240

35. Russek EM, Momennejad I, Botvinick MM, Gershman SJ, Daw ND (2017) Predictive representations can link model-based reinforcement learning to model-free mechanisms. PLoS Comput Biol 13:e1005768

36. Recanatesi S, Farrell M, Lajoie G, Deneve S, Rigotti M, Shea-Brown E (2021) Predictive learning as a network mechanism for extracting low-dimensional latent space representations. Nat Commun 12:1417

37. Daw ND, Niv Y, Dayan P (2005) Uncertainty-based competition between prefrontal and dorsolateral striatal systems for behavioral control. Nat Neurosci 8:1704–1711

38. Yu AJ, Dayan P (2005) Uncertainty, Neuromodulation, and Attention. Neuron 46:681–692

39. Broadbent DE, FitzGerald P, Broadbent MHP (1986) Implicit and explicit knowledge in the control of complex systems. British Journal of Psychology 77:33–50

40. Gonzalez C, Lerch JF, Lebiere C (2003) Instance-based learning in dynamic decision making. Cogn Sci 27:591–635

41. Bubic (2010) Prediction, cognition and the brain. Front Hum Neurosci. 10.3389/fnhum.2010.00025

42. Makarov VA, Villacorta-Atienza JA (2011) Compact Internal Representation as a Functional Basis for Protocognitive Exploration of Dynamic Environments. Recurrent Neural Networks for Temporal Data Processing. 10.5772/15127

43. Villacorta-Atienza JA, Calvo C, Makarov VA (2015) Prediction-for-CompAction: navigation in social environments using generalized cognitive maps. Biol Cybern 109:307–320

44. Diez-Hermano S, Aparicio-Rodriguez G, Manubens P, Sanchez-Jimenez A, Calvo-Tapia C, Levcik D, Villacorta-Atienza JA (2025) Minimal Neural Network Conditions for Encoding Future Interactions. Int J Neural Syst. 10.1142/S0129065725500169

45. Jezek K, Henriksen EJ, Treves A, Moser EI, Moser M-B (2011) Theta-paced flickering between place-cell maps in the hippocampus. Nature 478:246–249

46. Gupta AS, van der Meer MAA, Touretzky DS, Redish AD (2012) Segmentation of spatial experience by hippocampal theta sequences. Nat Neurosci 15:1032–1039

47. Lisman J, Redish AD (2009) Prediction, sequences and the hippocampus. Philosophical Transactions of the Royal Society B: Biological Sciences 364:1193– 1201

48. Skaggs WE, McNaughton BL, Wilson MA, Barnes CA (1996) Theta phase precession in hippocampal neuronal populations and the compression of temporal sequences. Hippocampus 6:149–172

49. Kim H, Schlichting ML, Preston AR, Lewis-Peacock JA (2020) Predictability Changes What We Remember in Familiar Temporal Contexts. J Cogn Neurosci 32:124–140

50. Schapiro AC, Kustner LV, Turk-Browne NB (2012) Shaping of Object Representations in the Human Medial Temporal Lobe Based on Temporal Regularities. Current Biology 22:1622–1627

51. Davachi L, DuBrow S (2015) How the hippocampus preserves order: the role of prediction and context. Trends Cogn Sci 19:92–99

52. Byrnes JP, Miller DC, Schafer WD (1999) Gender differences in risk taking: A meta-analysis. Psychol Bull 125:367–383

53. Harris CR, Jenkins M (2006) Gender Differences in Risk Assessment: Why do Women Take Fewer Risks than Men? Judgm Decis Mak 1:48–63

54. Ring P, Neyse L, David-Barett T, Schmidt U (2016) Gender Differences in Performance Predictions: Evidence from the Cognitive Reflection Test. Front Psychol. 10.3389/fpsyg.2016.01680

55. Hyde JS (2014) Gender Similarities and Differences. Annu Rev Psychol 65:373–398

56. Wraga M, Helt M, Jacobs E, Sullivan K (2007) Neural basis of stereotype-induced shifts in women’s mental rotation performance. Soc Cogn Affect Neurosci 2:12–19

57. Carothers BJ, Reis HT (2013) Men and women are from Earth: Examining the latent structure of gender. J Pers Soc Psychol 104:385–407

