## Supplementary material for "Event-Centered Prediction: How Future Interaction Points Shape Human Anticipation of Motion"

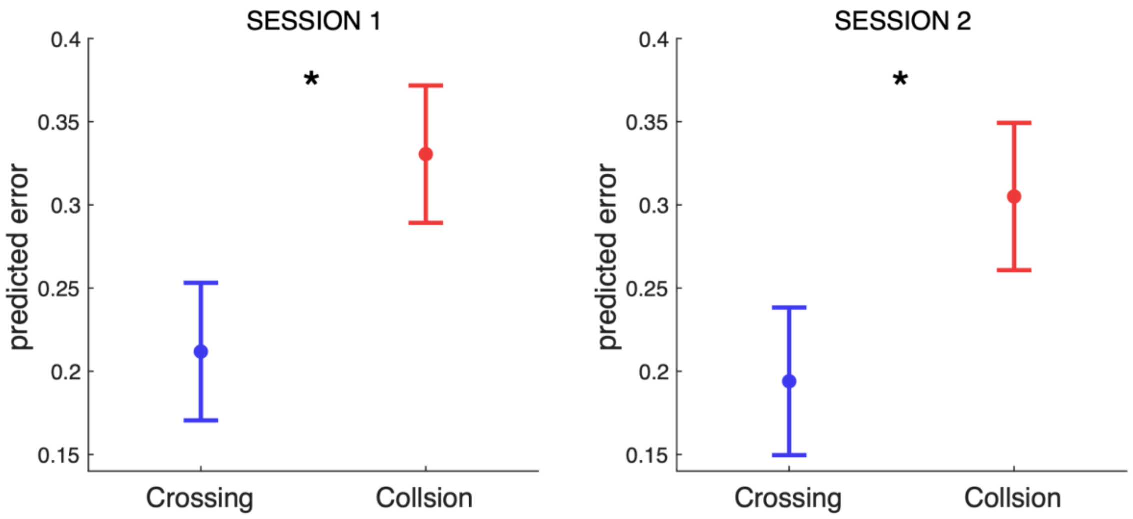


**Figure S1. Error in prediction in session 1 and 2.** Points and error bars represent model-based probability estimates and standard errors. Asterisks show significant differences between crossing and collision stimuli.

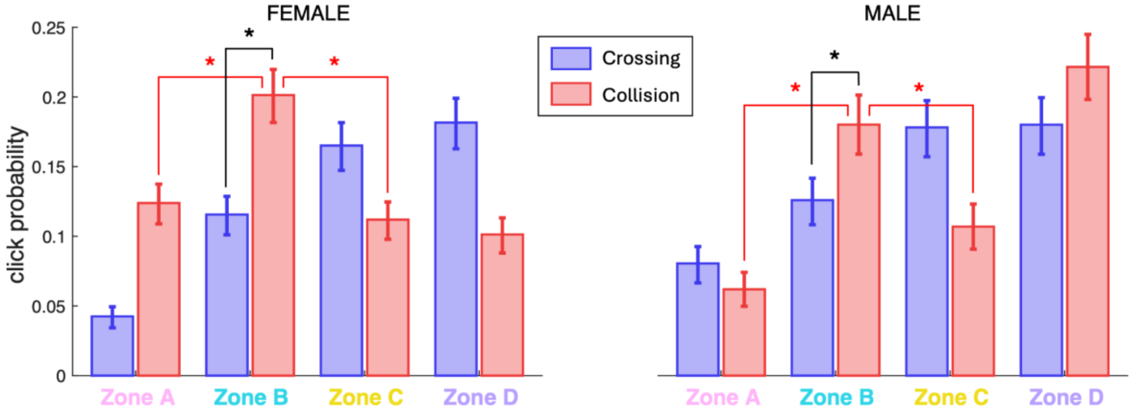


**Figure S2. Probability of click on each zone for Session 1.** Columns represent model-based probability estimates while error bars represent standard errors. Black asterisks show significant differences in Zone B between crossing and collision stimuli. Red asterisks show significant differences for collision stimuli between Zone B and its adjacent zones. Labels for each zone keep the same coloring as used in Figure 2.

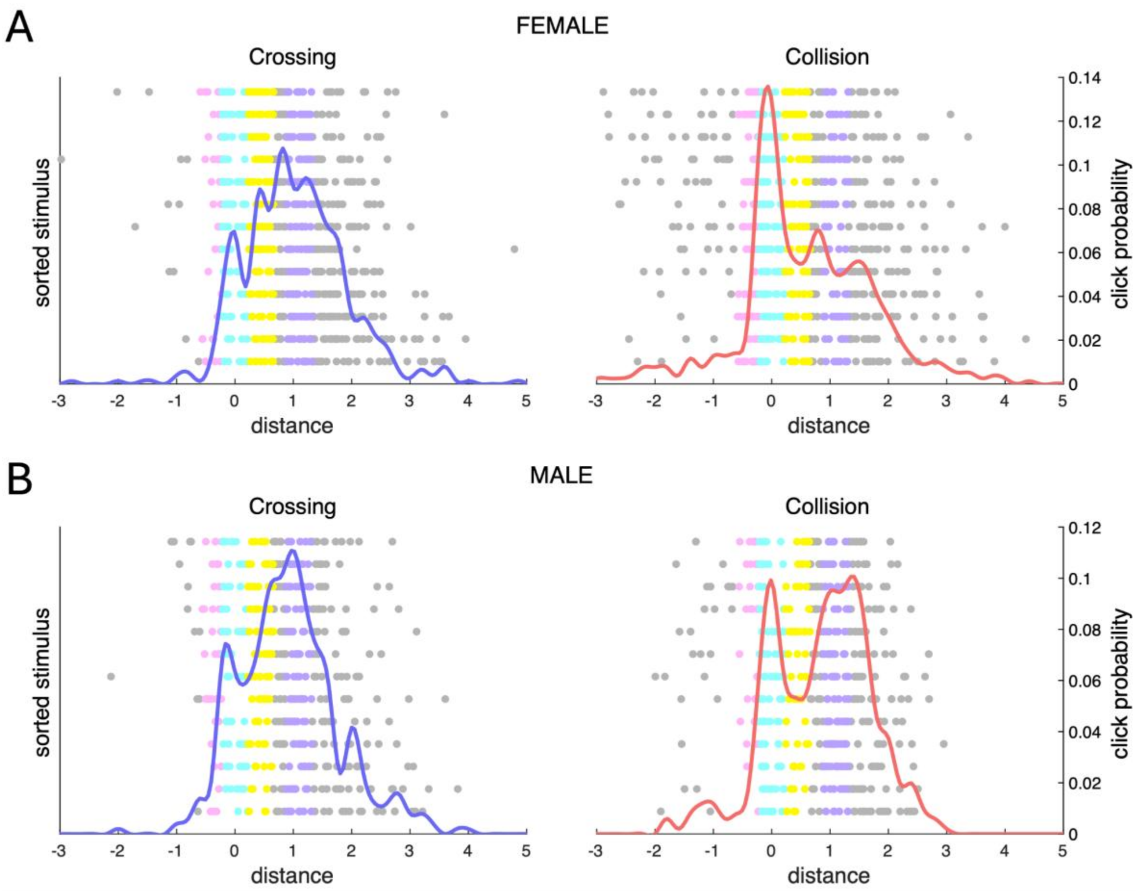


**Figure S3. Response patterns to each stimulus for Session 1. A.** Response patterns to each stimulus for female participants. Each dot represents a single click, plotted by distance (x-axis) and stimulus (y-axis). Stimuli are ordered by their mean click distance for collision stimuli, with the shortest mean distance at the top. Dot color indicates the corresponding zone (Fig. 2), while grey dots indicate clicks outside defined zones. The left panel shows cross trials (blue), and the right panel shows collision trials (red), with the corresponding aggregate distributions, represented by the colored function. **B.** Response patterns to each stimulus for male participants. Same conventions as in A.

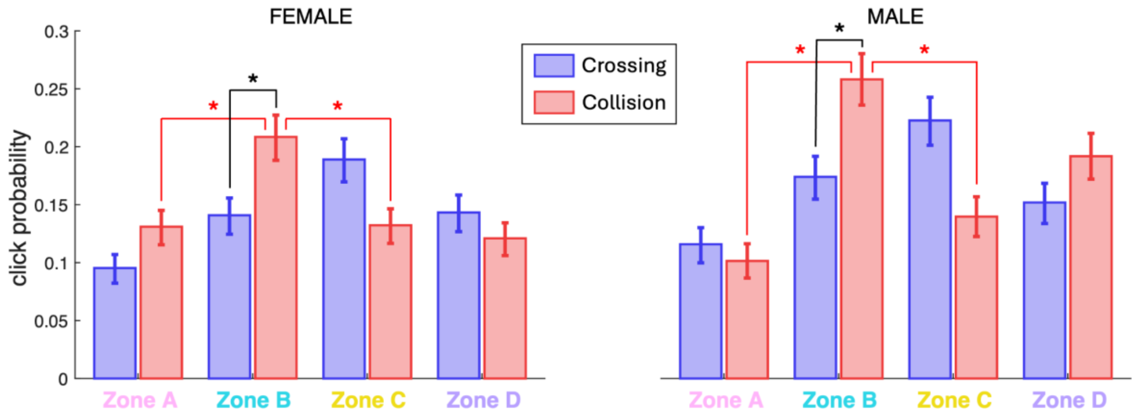


**Figure S4. Probability of click on each zone for Session 2.** Columns represent model-based probability estimates while error bars represent standard errors. Black asterisks show significant differences in Zone B between crossing and collision stimuli. Red asterisks show significant differences for collision stimuli between Zone B and its adjacent zones. Labels for each zone keep the same coloring as used in Figure 2.

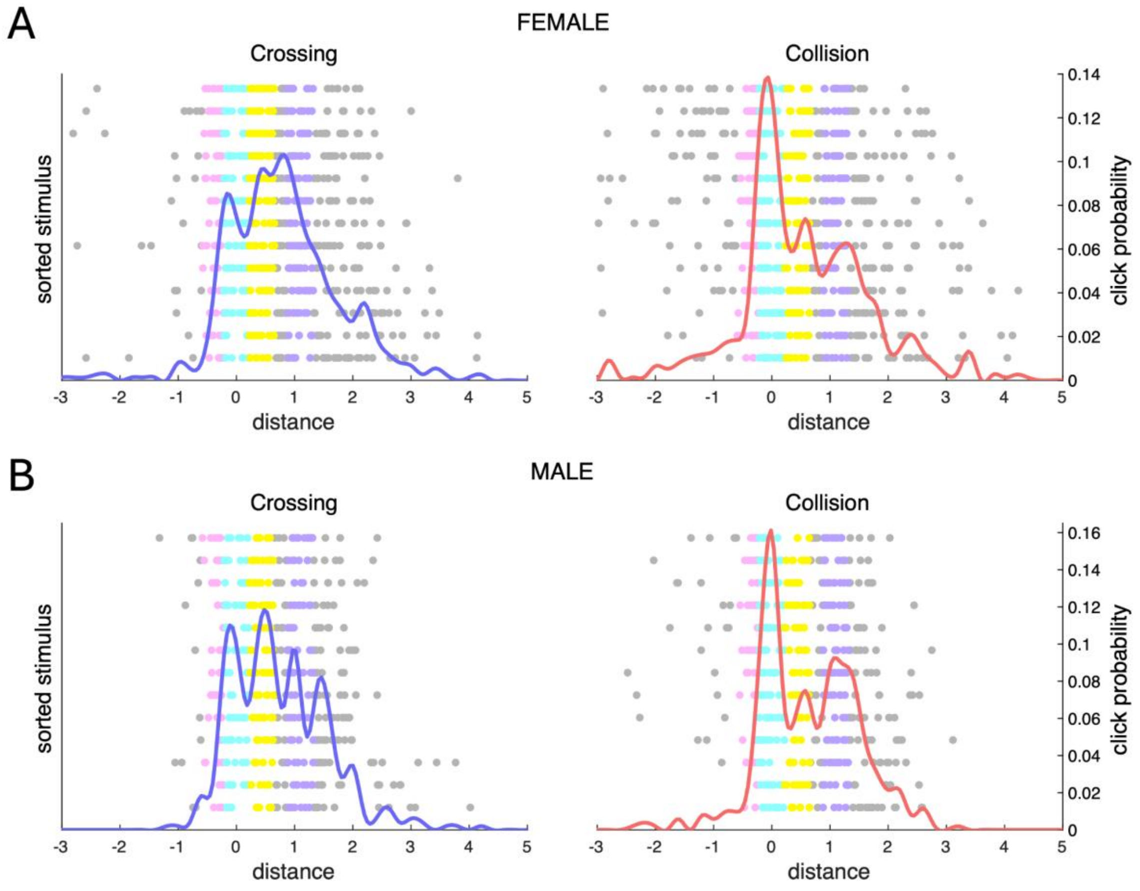


**Figure S5. Response patterns to each stimulus for Session 2. A.** Response patterns to each stimulus for female participants. Each dot represents a single click, plotted by distance (x-axis) and stimulus (y-axis). Stimuli are ordered by their mean click distance for collision stimuli, with the shortest mean distance at the top. Dot color indicates the corresponding zone (Fig. 2), while grey dots indicate clicks outside defined zones. The left panel shows cross trials (blue), and the right panel shows collision trials (red), with the corresponding aggregate distributions, represented by the colored function. **B.** Response patterns to each stimulus for male participants. Same conventions as in A.

Table S1. Frequencies and age description of the participants.

| Gender | Absolute frequency | Relative frequency | Mean age | Median age | Quantile 0.05 for age | Quantile 0.95 for age |
|---|---|---|---|---|---|---|
| Female | 64 | 0.615 | 19.9 | 18.5 | 18 | 24.8 |
| Male | 40 | 0.385 | 20 | 18.5 | 18 | 25 |

Table S2. Session 1: transformed error mixed model

| Predictor | Estimate | CI* | p-value |
|---|---|---|---|
| (Intercept) | 0.20 | 0.06 – 0.35 | **0.005** |

| | | | |
|---|---|---|---|
| type [Cross] | -0.39 | -0.54 – -0.23 | **<0.001** |

$\tau_{00}$ id = 0.24 (n = 104); $\tau_{00}$ stimulus = 0.03 (n = 26); ICC = 0.28; 2548 observations

*α = 0.05

Table S3. Session 2: transformed error mixed model

| Predictor | Estimate | CI* | p-value |
|---|---|---|---|
| (Intercept) | 0.18 | 0.05 – 0.32 | **0.008** |
| type [Cross] | -0.35 | -0.47 – -0.23 | **<0.001** |

$\tau_{00}$ id = 0.29 (n = 104); $\tau_{00}$ stimulus = 0.02 (n = 26); ICC = 0.32; 2502 observations

*α = 0.05

Table S4. Error comparisons among sessions: transformed error mixed model

| Predictor | Estimate | CI* | p-value |
|---|---|---|---|
| (Intercept) | 0.25 | 0.11 – 0.38 | **<0.001** |
| type [Cross] | -0.37 | -0.50 – -0.24 | **<0.001** |
| session [2] | -0.11 | -0.15 – -0.06 | **<0.001** |

$\tau_{00}$ id = 0.25 (n = 104); $\tau_{00}$ stimulus = 0.02 (n = 26); ICC = 0.27; 5050 observations

*α = 0.05

Table S5. Error comparisons among sessions: untransformed error mixed model

| Predictor | Estimate | CI* | p-value |
|---|---|---|---|
| (Intercept) | 0.33 | 0.29 – 0.37 | **<0.001** |
| type [Cross] | -0.12 | -0.15 – -0.09 | **<0.001** |
| session [2] | -0.02 | -0.04 – -0.01 | **0.003** |

$\tau_{00}$ id = 0.03 (n = 104); $\tau_{00}$ stimulus > 0.01 (n = 26); ICC = 0.31; 5050 observations

*α = 0.05

Table S6. Session 1: click distribution comparisons (two_sample test)

| Comparison | Statistic | p-value |
|---|---|---|
| Men collision vs men cross | 15.022 | **0.002** |
| Women collision vs women cross | 39.110 | **<0.001** |
| Men collision vs women collision | 29.811 | **<0.001** |
| Men cross vs women cross | 11.935 | 0.107 |

Table S7. Session 1: triple interaction area clicking model for collisions (Anova comparisons on table below)

| Predictor | Estimate | OR | OR CI* | p-value |
|---|---|---|---|---|
| (Intercept) | -1.37 | 0.25 | 0.20 – 0.32 | **<0.001** |
| gender [Male] | -0.16 | 0.85 | 0.59 – 1.22 | 0.388 |
| type [Cross] | -0.65 | 0.52 | 0.40 – 0.68 | **<0.001** |
| area [areaA] | -0.58 | 0.56 | 0.43 – 0.73 | **<0.001** |
| area [areaC] | -0.69 | 0.50 | 0.38 – 0.66 | **<0.001** |
| area [areaD] | -0.80 | 0.45 | 0.34 – 0.59 | **<0.001** |
| Gender [Male] x type [Cross] | 0.23 | 1.26 | 0.81 – 1.95 | 0.301 |
| Gender [Male] x area [areaA] | -0.63 | 0.53 | 0.33 – 0.87 | **0.012** |
| Gender [Male] x area [areaC] | 0.08 | 1.09 | 0.69 – 1.70 | 0.719 |
| Gender [Male] x area [areaD] | 1.06 | 2.89 | 1.91 – 4.39 | **<0.001** |
| type [Cross] × area [areaA] | -0.50 | 0.61 | 0.38 – 0.97 | **0.037** |
| type [Cross] × area [areaC] | 1.10 | 3.02 | 2.04 – 4.47 | **<0.001** |
| type [Cross] × area [areaD] | 1.33 | 3.78 | 2.55 – 5.62 | **<0.001** |

| | | | | |
|---|---|---|---|---|
| (gender [Male] × type [Cross]) × area [areaA] | 1.21 | 3.35 | 1.59 – 7.05 | **0.001** |
| (gender [Male] × type [Cross]) × area [areaC] | -0.09 | 0.92 | 0.49 – 1.72 | 0.790 |
| (gender [Male] × type [Cross]) × area [areaD] | -1.17 | 0.31 | 0.17 – 0.57 | **<0.001** |

$\tau_{00}$ id = 0. 32 (n = 104); ICC = 0.09; 10192 observations

*α = 0.05

Table S8. Session 1: Anova comparisons for triple interaction area clicking model from S7

| Predictor | Chisq | Df | p-value |
|---|---|---|---|
| Gender | 0.432 | 1 | 0.511 |
| Type | 0.448 | 1 | 0.503 |
| Area | 89.469 | 3 | **<0.001** |
| Gender:type | 0.058 | 1 | 0.809 |
| Gender:area | 13.549 | 3 | **0.004** |
| Type:area | 65.885 | 3 | **<0.001** |
| Gender:type:area | 42.356 | 3 | **<0.001** |

Table S9. Session 1: area clicking model for women

| Predictor | Estimate | OR | OR CI* | p-value |
|---|---|---|---|---|
| (Intercept) | -1.38 | 0.25 | 0.20 – 0.32 | **<0.001** |
| type [Cross] | -0.66 | 0.52 | 0.39 – 0.68 | **<0.001** |
| areas [areaA] | -0.58 | 0.56 | 0.43 – 0.73 | **<0.001** |

| | | | | |
|---|---|---|---|---|
| areas [areaC] | -0.69 | 0.50 | 0.38 – 0.66 | **<0.001** |
| areas [areaD] | -0.80 | 0.45 | 0.34 – 0.59 | **<0.001** |
| type [Cross] × areas [areaA] | -0.50 | 0.61 | 0.38 – 0.97 | **0.037** |
| type [Cross] × areas [AreaC] | 1.11 | 3.03 | 2.04 – 4.49 | **<0.001** |
| type [Cross] × areas [AreaD] | 1.33 | 3.80 | 2.56 – 5.64 | **<0.001** |

$\tau_{00}$ id = 0. 39 (n = 64);  ICC = 0.11; 6148 observations

*α = 0.05

Table S10. Session 1: comparisons between types among women by area

| Area | Estimate | SE | p-value |
|---|---|---|---|
| Area A | 1.16 | 0.20 | **<0.001** |
| Area B | 0.66 | 0.14 | **<0.001** |
| Area C | -0.45 | 0.14 | **0.001** |
| Area D | -0.68 | 0.15 | **<0.001** |

Contrasts shown are Collision – Cross

Table S11. Session 1: comparisons between area B and the other areas among women

| Type | Area | Estimate | SE | p-value |
|---|---|---|---|---|
| Collision | Area A | 0.58 | 0.14 | **<0.001** |
| | Area C | 0.69 | 0.14 | **<0.001** |
| | Area D | 0.81 | 0.14 | **<0.001** |
| Cross | Area A | 1.08 | 0.20 | **<0.001** |
| | Area C | -0.41 | 0.14 | **0.021** |
| | Area D | -0.53 | 0.14 | **0.001** |

Contrasts shown are area B – other areas

Table S12. Session 1: area clicking model for men

| Predictor | Estimate | OR | OR CI* | p-value |
|---|---|---|---|---|
| (Intercept) | -1.52 | 0.22 | 0.17 – 0.29 | **<0.001** |
| type [Cross] | -0.42 | 0.66 | 0.47 – 0.92 | **0.015** |
| areas [areaA] | -1.20 | 0.30 | 0.20 – 0.45 | **<0.001** |
| areas [areaC] | -0.61 | 0.54 | 0.38 – 0.78 | **0.001** |
| areas [areaD] | 0.26 | 1.30 | 0.95 – 1.76 | 0.099 |
| type [Cross] × areas [areaA] | 0.70 | 2.02 | 1.14 – 3.60 | **0.017** |
| type [Cross] × areas [AreaC] | 1.02 | 2.76 | 1.69 – 4.52 | **<0.001** |
| type [Cross] × areas [AreaD] | 0.16 | 1.18 | 0.74 – 1.86 | 0.486 |

$\tau_{00}$ id = 0. 23 (n = 40); ICC = 0.06; 4044 observations

*α = 0.05

Table S13. Session 1: comparisons between types among men by area

| Area | Estimate | SE | p-value |
|---|---|---|---|
| Area A | -0.28 | 0.24 | 0.235 |
| Area B | 0.42 | 0.17 | **0.015** |
| Area C | -0.59 | 0.18 | **<0.001** |
| Area D | 0.26 | 0.16 | 0.098 |

Contrasts shown are Collision – Cross

Table S14. Session 1: comparisons between area B and the other areas among men

| Type | Area | Estimate | SE | p-value |
|---|---|---|---|---|
| Collision | Area A | 1.20 | 0.21 | **<0.001** |
| | Area C | 0.61 | 0.18 | **0.004** |
| | Area D | -0.25 | 0.16 | 0.350 |

| | | | | |
|---|---|---|---|---|
| Cross | Area A | 0.498 | 0.21 | 0.073 |
| | Area C | -0.41 | 0.17 | 0.089 |
| | Area D | -0.00 | 0.16 | >0.999 |

Contrasts shown are area B – other areas

Table S15. Session 1 vs Session 2: click distribution comparisons (two_sample test)

| Group | Statistic | p-value |
|---|---|---|
| Men collision | 13.203 | **0.003** |
| Men cross | 12.208 | **0.007** |
| Women collision | 7.924 | 0.661 |
| Women cross | 21.627 | **<0.001** |

Table S16. Session 2: triple interaction area clicking model for collisions (Anova comparisons on table below)

| Predictor | Estimate | OR | OR CI* | p-value |
|---|---|---|---|---|
| (Intercept) | -1.32 | 0.27 | 0.22 – 0.33 | **<0.001** |
| gender [Men] | 0.24 | 1.27 | 0.91 – 1.78 | 0.153 |
| type [Cross] | -0.47 | 0.62 | 0.48 – 0.81 | **<0.001** |
| area [areaA] | -0.56 | 0.57 | 0.44 – 0.75 | **<0.001** |
| area [areaC] | -0.55 | 0.58 | 0.44 – 0.76 | **<0.001** |
| area [areaD] | -0.65 | 0.52 | 0.40 – 0.69 | **<0.001** |
| Gender [Men] x type [Cross] | -0.03 | 0.97 | 0.64 – 1.45 | 0.865 |
| Gender [Men] x area [areaA] | -0.58 | 0.56 | 0.36 – 0.87 | **0.009** |
| Gender [Men] x area [areaC] | -0.22 | 0.80 | 0.53 – 1.21 | 0.290 |
| Gender [Men] x area [areaD] | 0.26 | 1.30 | 0.87 – 1.94 | 0.206 |
| type [Cross] × area | 0.12 | 1.12 | 0.75 – 1.69 | 0.574 |

| | | | | |
|---|---|---|---|---|
| [areaA] | | | | |
| type [Cross] × area [areaC] | 0.90 | 2.45 | 1.68 – 3.58 | **<0.001** |
| type [Cross] × area [areaD] | 0.67 | 1.95 | 1.32 – 2.88 | **0.001** |
| (gender [Men] × type [Cross]) × area [areaA] | 0.54 | 1.71 | 0.90 – 3.26 | 0.100 |
| (gender [Men] × type [Cross]) × area [areaC] | 0.18 | 1.20 | 0.67 – 2.16 | 0.541 |
| (gender [Men] × type [Cross]) × area [areaD] | -0.44 | 0.64 | 0.35 – 1.16 | 0.144 |

$\tau_{00}$ id = 0. 26 (n = 104); ICC = 0.07; 10008 observations

*α = 0.05

Table S17. Session 2: Anova comparisons for triple interaction area clicking model

| Predictor | Chisq | Df | p-value |
|---|---|---|---|
| Gender | 1.51 | 1 | 0.219 |
| Type | 0.73 | 1 | 0.392 |
| Area | 68.72 | 3 | **<0.001** |
| Gender:type | 0.003 | 1 | 0.959 |
| Gender:area | 5.53 | 3 | 0.137 |
| Type:area | 44.42 | 3 | **<0.001** |
| Gender:type:area | 9.01 | 3 | **0.029** |

Table S18. Session 2: area clicking model for women

| Predictor | Estimate | OR | CI* | p-value |
|---|---|---|---|---|
| (Intercept) | -1.33 | 0.26 | 0.21 – 0.33 | **<0.001** |
| type [Cross] | -0.48 | 0.62 | 0.48 – 0.81 | **<0.001** |
| areas [areaA] | -0.56 | 0.57 | 0.44 – 0.75 | **<0.001** |
| areas [areaC] | -0.55 | 0.58 | 0.44 – 0.75 | **<0.001** |
| areas [areaD] | -0.65 | 0.52 | 0.40 – 0.69 | **<0.001** |
| type [Cross] × areas [areasA] | 0.12 | 1.13 | 0.75 – 1.69 | 0.571 |
| type [Cross] × areas [AreaC] | 0.90 | 2.46 | 1.68 – 3.60 | **<0.001** |
| type [Cross] × areas [AreaD] | 0.67 | 1.95 | 1.32 – 2.89 | **0.001** |

$\tau_{00}$ id = 0. 38 (n = 64); ICC = 0.10; 5992 observations

*α = 0.05

Table S19. Session 2: comparisons between types among women by area

| Area | Estimate | SE | p-value |
|---|---|---|---|
| Area A | 0.36 | 0.16 | **0.024** |
| Area B | 0.48 | 0.14 | **<0.001** |
| Area C | -0.43 | 0.14 | **0.002** |
| Area D | -0.20 | 0.15 | 0.187 |

Contrasts shown are Collision – Cross

Table S20. Session 2: comparisons between area B and the other areas among women

| Type | Area | Estimate | SE | p-value |
|---|---|---|---|---|
| Collision | Area A | 0.56 | 0.14 | **<0.001** |
| | Area C | 0.55 | 0.14 | **<0.001** |
| | Area D | 0.65 | 0.14 | **<0.001** |

| | | | | |
|---|---|---|---|---|
| Cross | Area A | 0.44 | 0.16 | **0.026** |
| | Area C | -0.35 | 0.14 | 0.052 |
| | Area D | -0.02 | 0.14 | 0.999 |

Contrasts shown are area B – other areas

Table S21. Session 2: area clicking model for men

| Predictor | Estimate | OR | CI* | p-value |
|---|---|---|---|---|
| (Intercept) | -1.06 | 0.35 | 0.28 – 0.43 | **<0.001** |
| type [Cross] | -0.50 | 0.61 | 0.45 – 0.82 | **0.001** |
| areas [areaA] | -1.12 | 0.32 | 0.23 – 0.46 | **<0.001** |
| areas [areaC] | -0.76 | 0.47 | 0.34 – 0.64 | **<0.001** |
| areas [areaD] | -0.38 | 0.68 | 0.51 – 0.92 | **0.011** |
| type [Cross] × areas [areaA] | 0.65 | 1.92 | 1.17 – 3.15 | **0.010** |
| type [Cross] × areas [AreaC] | 1.07 | 2.91 | 1.87 – 4.55 | **<0.001** |
| type [Cross] × areas [AreaD] | 0.22 | 1.25 | 0.80 – 1.95 | 0.335 |

$\tau_{00}$ id = 0. 15(n = 40); ICC = 0.04; 4016 observations

*α = 0.05

Table S22. Session 2: comparisons between types among men by area

| Area | Estimate | SE | p-value |
|---|---|---|---|
| Area A | -0.15 | 0.20 | 0.457 |
| Area B | 0.50 | 0.16 | **0.001** |
| Area C | -0.57 | 0.17 | **0.001** |
| Area D | 0.28 | 0.17 | 0.092 |

Contrasts shown are Collision – Cross

Table S23. Session 2: comparisons between area B and the other areas among men

| Type | Area | Estimate | SE | p-value |
|---|---|---|---|---|
| Collision | Area A | 1.13 | 0.18 | **<0.001** |
| | Area C | 0.76 | 0.16 | **<0.001** |
| | Area D | 0.38 | 0.15 | 0.053 |
| Cross | Area A | 0.48 | 0.18 | **0.045** |
| | Area C | -0.31 | 0.16 | 0.221 |
| | Area D | 0.16 | 0.17 | 0.779 |

Contrasts shown are area B – other areas
